# The fully resolved genome of *Bacillus thuringiensis* HER1410 reveals a *cry*-containing chromosome, two megaplasmids & an integrative plasmidial prophage

**DOI:** 10.1101/2020.05.05.080028

**Authors:** Ana Lechuga, Cédric Lood, Margarita Salas, V. Vera van Noort, Rob Lavigne, Modesto Redrejo-Rodríguez

## Abstract

*Bacillus thuringiensis* is the most used biopesticide in agriculture. Its entomopathogenic capacity stems from the possession of plasmid-borne insecticidal crystal genes (*cry*), traditionally used as discriminant taxonomic feature for that species. As such, crystal and plasmid identification are key to the characterization of this species. To date, about 600 *B. thuringiensis* genomes have been reported, but less than 5% have been resolved, while the other draft genomes are incomplete, precluding plasmid delineation. Here we present the complete genome of *Bacillus thuringiensis* HER1410, a strain closely related to *B. thuringiensis entomocidus* and a known host for a variety of *Bacillus* phages. The combination of short and long-reads techniques allowed fully resolving the genome and delineation of three plasmids. This enabled the accurate detection of an unusual location of a unique *cry* gene, *cry1Ba4,* located in a genomic island near the chromosome replication origin. Two megaplasmids, pLUSID1 and pLUSID2 could be delineated: pLUSID1 (368kb), a likely conjugative plasmid involved in virulence, and pLUSID2 potentially related to the sporulation process. A smaller plasmidial prophage pLUSID3, with a dual lifestyle whose integration within the chromosome, causes the disruption of a flagellar key component. Finally, phylogenetic analysis located this strain within a clade comprising members from the *B. thuringiensis* serovar *thuringiensis* and other serovars and with *B.cereus s. s.* This highlights the intermingled taxonomy of *B. cereus sensu lato* group, where genomics alone does not support the present taxonomy between *B. cereus s. s.* and *B. thuringiensis* as species designation currently relies solely on the presence of entomocidal genes.

**Importance:** *Bacillus cereus* group species have been extensively studied due to their economical and clinical relevance. This importance originally set the basis for *B. cereus* group members classification which are commonly based on phenotypical criteria. Sequencing era has shed light about genomic characterization of these species, showing their chromosomal genomic similarity and highlighting the role of mobile genetic elements, especially megaplasmids, in the classification and characterization of this group. However, only the 5% of the sequenced *B. thuringiensis* genomes have been fully resolved. Thus, here we addressed efficiently the study *B. thuringiensis* HER1410 genomic features by the use of a combination of short and long-reads sequencing. This methodology resulted in the high-quality assembly, which led to the identification of an uncommon location of a *cry* gene close to the chromosomal origin, as well as three fully resolved extrachromosomal elements, two megaplasmids and an integrative plasmidial prophage.

## Introduction

The *Bacillus cereus* group encompasses 21 published species with a common monophyletic origin that display highly differentiated phenotypes in terms of ecological niches and virulence spectra (Liu et al., 2017, Carroll et al., 2020). Among them, some have been extensively studied due to their medical and economic importance. This group contains well-known pathogenic species including *Bacillus cereus sensu stricto*, linked to food poisoning, emesis and diarrhea, and *Bacillus anthracis*, the etiological agent of anthrax. There are also non-pathogenic industrially relevant species like *Bacillus toyonensis*, used as probiotic in veterinary medicine and *Bacillus thuringiensis,* which produces insecticidal toxins and is often used as an industrial biopesticide control agent (Ehling-Schulz et al., 2019). These phenotypes often find their genetic origin on large plasmids where toxins such as anthrax, bioinsecticidal crystal proteins, or emetic toxins are encoded. Toxin-bearing (mega)plasmids, as well as a variety of smaller plasmids, comprise the vast diversity of accessory elements characteristic of the *B. cereus* group. This accessory genome, as well as other mobile genetic elements (MGEs) like prophages and genomic islands, favor a rapid adaptative response to ecological change (Raymond and Bonsall, 2013, Fu et al., 2019).

Traditionally, *B. cereus* group members have been classified based on phenotypical criteria associated with their clinical and industrial relevance. However, these defined phenotypes do not necessarily agree with a genomic-based taxonomy as the plasmids causing the phenotypes can be transferred between less related species. Several recent studies addressing this problem are suggesting a new taxonomic nomenclature for this group (Baek et al., 2019, Carroll et al., 2020). A clear example of this problem is the extensive sequence similarity between *B. cereus s. s*. and *B. thuringiensi*s. The latter is commonly discriminated based on the occurrence of parasporal crystal inclusion bodies. In some cases, this phenotypical classification can lead to an inaccurate labelling as *B. thuringiensis* of some *B. cereus* isolates that are capable of causing both diarrheal foodborne disease and producing bioinsecticidal crystal proteins (Johler et al., 2018). Two types of proteins, Cry (crystal) and Cyt (cytolytic) proteins, are the principal components of these crystal protein inclusions and have shown high specific activity against different insects (Ehling-Schulz et al., 2019). *Cry* and *cyt* genes are located in plasmids such as the well-studied *B. thuringiensis* ser. *israelensis* pBtoxis which harbors up to five different *cry* genes and three *cyt* genes (Gillis et al., 2018). These genes are usually flanked by mobile elements (insertion sequences and transposons) and can grouped with other insecticidal proteins to form pathogenicity islands (PAI). This organization and the surrounding genetic context of these genes have been described to facilitate their mobility among plasmids and strains (Fiedoruk et al., 2017).

Numerous tailed bacteriophages preying *Bacillus cereus sensu lato* have been previously identified and, among them, a recently accepted genus of the *Tectiviridae* family, *Betatectivirus* was created (Gillis and Mahillon, 2014a). The recently renewed interest in the study of tectiviruses stems from the narrow host specificity of some members for dangerous human pathogens, their complex diversity and variability patterns, and a possible phylogenetic relationship with some groups of eukaryotic viruses and mobile elements (Krupovic and Koonin, 2015). In the study of betatectiviruses, *B. thuringiensis* GBJ002, HER1410, and *B. cereus* HER1047 strains have been used as hosts for tectiviral infections. These three strains represent different serotypes and have been especially useful to study tectiviruses features such as virus entry and host cell physiology (Gaidelyte et al., 2006, Daugelavicius et al., 2007). However, despite their importance to this field of research, none of them have been sequenced to date. In particular, *Bacillus thuringiensis* HER1410 is extensively used as a host of Bam35, the model species of *Betatectivirus,* for the molecular and structural studies on this virus (Ravantti et al., 2003, Laurinmaki et al., 2005, Berjon-Otero et al., 2016). Further, HER1410 has shown high sensitivity to different *Bacillus* phages families allowing, for instance, the identification of novel tectiviruses (Gillis and Mahillon, 2014b). Besides, HER1410 has been also included in *B. cereus* group studies on hemolytic activity and temperature dependent growth rates (Francis et al., 1998, Pruss et al., 1999a, Pruss et al., 1999b).

This study aims to characterize *Bacillus thuringiensis* HER1410, known as a sensitive host of a large variety of *Bacillus* phages and a reference strain to study tectiviruses. A combination of short and long-reads sequencing methods was applied, known to allow the full resolution of bacterial genomes and delineation of plasmids (Albers et al., 2018, T’Syen et al., 2018, Botelho et al., 2019). We successfully resolved the chromosome and identified and characterized three extrachromosomal elements comprising the HER1410 genome and present a detailed overview of these components, focusing on the key elements in *B. cereus* group such as the description of MGEs, virulence factors and phylogeny.

## Materials and methods

### BACTERIAL STRAIN

*Bacillus thuringiensis* HER1410 strain was from Laboratory Stock. The original strain was obtained from culture collection Félix d’Herelle Reference Center for Bacterial Viruses of the Université of Laval (https://www.phage.ulaval.ca).

### DNA EXTRACTION AND GENOME SEQUENCING

*B. thuringiensis* HER1410 was inoculated in LB medium and incubated overnight at 37° to an OD600 of 1.14. Genomic DNA (gDNA) was isolated using the DNeasy Blood and Tissue kit (Qiagen) and purified by ethanol precipitation. The quality of the gDNA was checked using a ThermoFisher Scientific NanoDrop spectrophotometer (OD280/260 and OD230/260), and its integrity was verified by gel electrophoresis (1% agarose w/v).

The gDNA was sequenced using Illumina and Nanopore technologies. The first set of reads was obtained on an Illumina MiniSeq using a paired-end 2×150 bp approach with a library obtained using Nextera Flex (Illumina, inc.). A second set of reads was obtained on a MinION nanopore sequencer (Oxford Nanopore Technology) equipped with a flowcell of type R9.4.1 and a library prepared using the 1D ligation approach with native barcoding (Oxford Nanopore Technology).

### GENOME HYBRID ASSEMBLY AND FUNCTIONAL ANNOTATION

The Illumina reads were controlled for quality using FastQC v0.11.8 and trimmomatic v0.38 for adapter clipping, quality trimming (LEADING:3 TRAILING:3 SLIDINGWINDOW:4.15), and minimum length exclusion (>50 bp) (Andrews, 2010, Bolger et al., 2014). The quality of the nanopore reads was evaluated visually using Nanoplot v1.27.0 (De Coster et al., 2018) and then further processed using Porechop v0.2.3 to clip barcodes and with NanoFilt v2.3.0 for quality exclusion (Q>10) and length (>1,000 bp) (Wick, 2017).

The genome was assembled *de novo* using the short-reads SPAdes assembler v3.13.1 with default options, and the hybrid Unicycler v0.4.8 assembler using bold mode (Bankevich et al., 2012, Wick et al., 2017). The assembly metrics reported in Supplemental Figure 1 were computed using QUAST v4 and the visualization of the assembly graphs was drawn with Bandage v0.8.1 (Gurevich et al., 2013, Wick et al., 2015).

**Figure 1.**
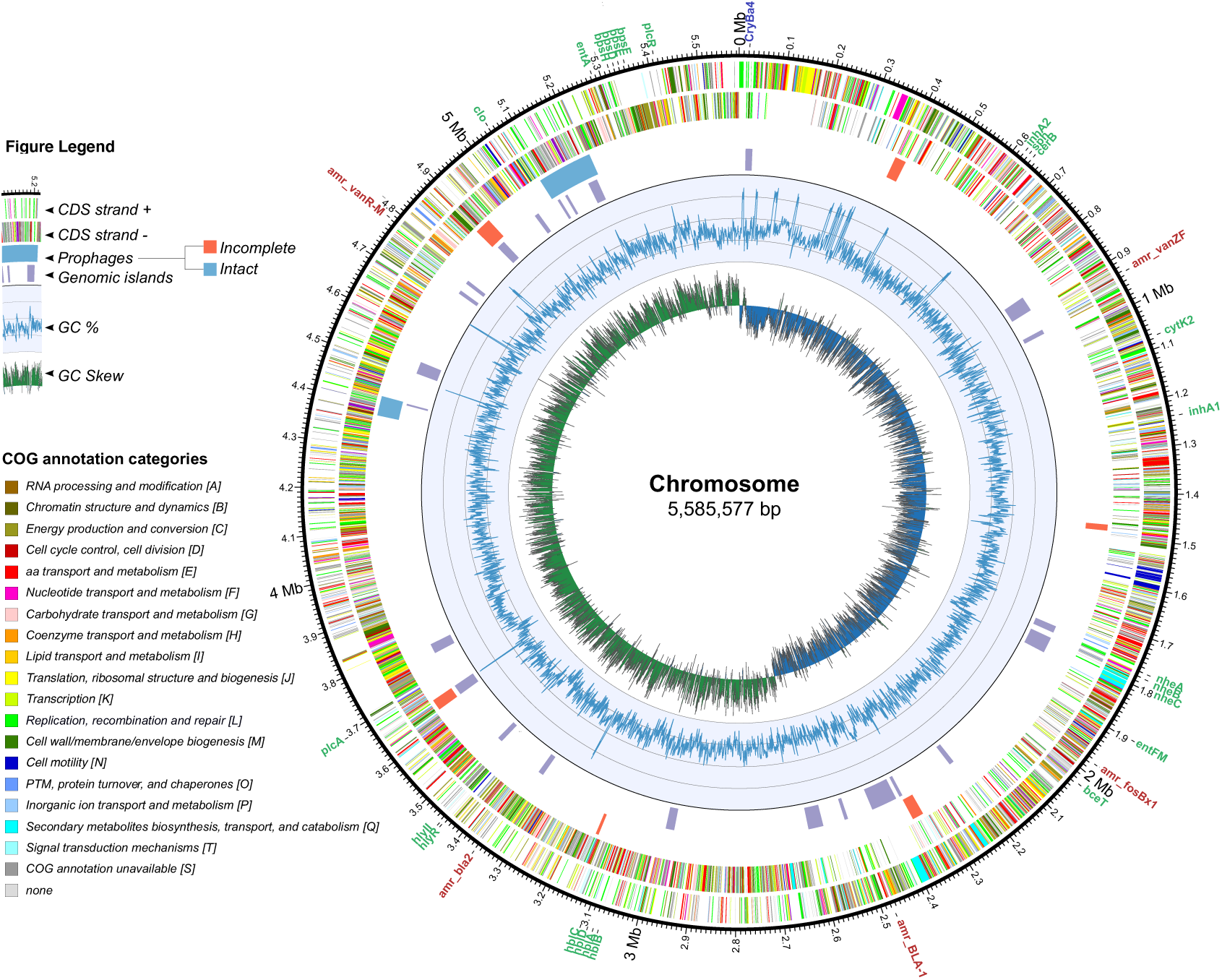
Circular representation of *B. thuringiensis* HER1410 chromosome. The outer labels refer to toxin (green) and antibiotic resistance genes (red), found using the bTyper and bTyper3 tools. The next two circles represent the coding regions on the positive and negative strands colored by their functional annotation. The next two circles indicate the genomic islands predicted by the IslandViewer package (purple) and prophages predicted by PHASTER webserver where orange=incomplete, blue=intact. The inner two rings indicate the changes in %GC content and the GC skew respectively. Coordinates of the highlighted features can be found in Supplemental Table 3. Chromosomal integration of pLUSID3 was excluded from the chromosome sequence and, therefore, this representation.

The annotation of the genome was performed using the NCBI Prokaryotic Genome Annotation Pipeline (PGAP), and the resulting annotated proteome was further annotated with COG categories and KEGG pathways using the eggNOG mapper (Tatusova et al., 2016, Huerta-Cepas et al., 2017). The annotation of the *cry1Ba4*-containing island was updated using ISFinder (Siguier, 2006) (Figure 2, Supplemental Table 3). The annotation of the extrachromosomal elements was updated using eggNOG mapper (Huerta-Cepas et al., 2017), PHASTER (Arndt et al., 2016) and PHMMER tools (Potter et al., 2018) (Figures 3 and 4, Supplemental tables 3 and 4).

**Figure 2.**
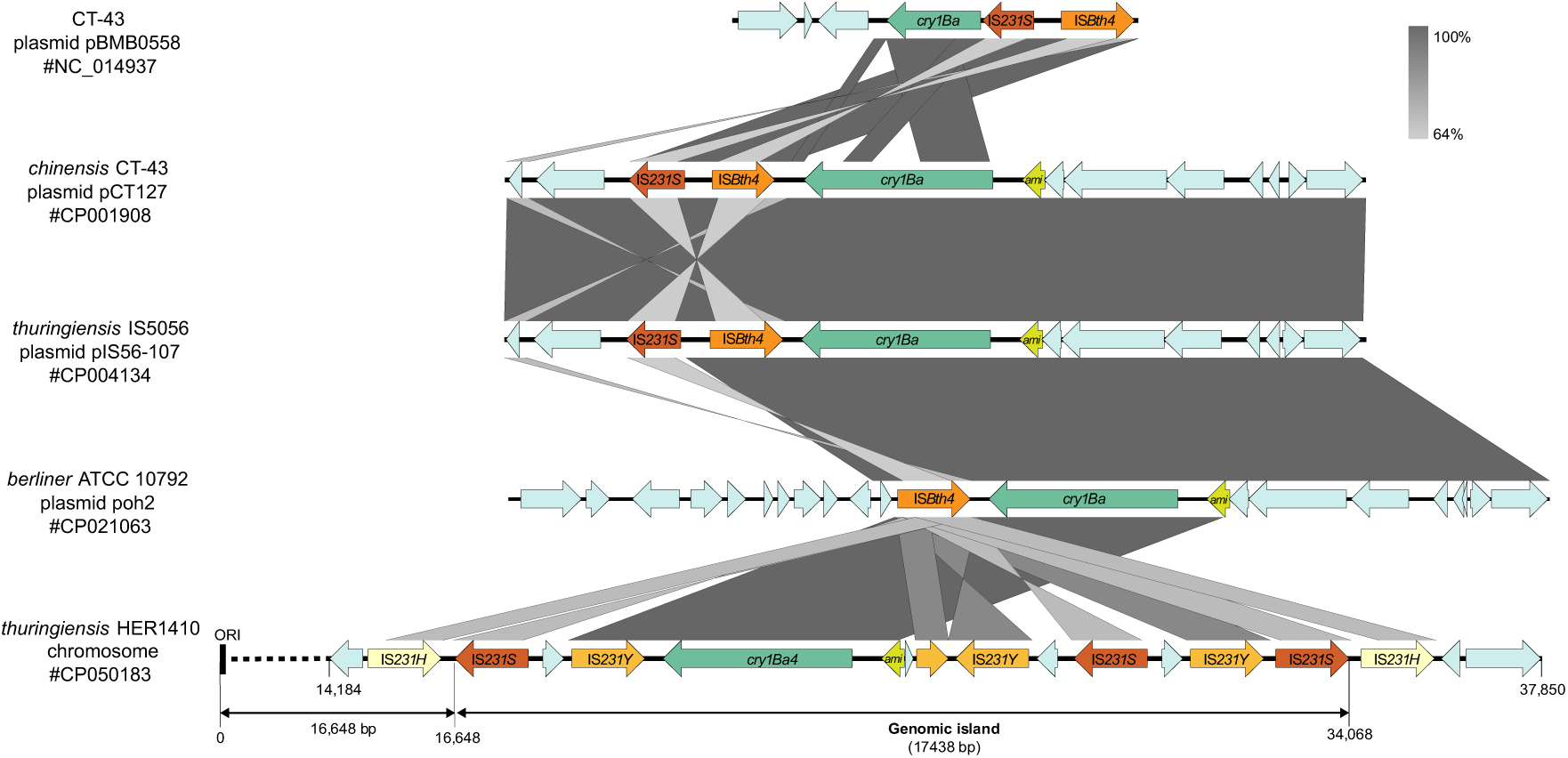
HER1410 chromosomal *cryBa4*-containing genomic island analysis. Comparison of *cry1Ba4*-containing genomic island with described *cry1Ba*-containing plasmid cassettes from Fiedoruk et al. (2017) was rendered with EasyFig and further details were added with Adobe Illustrator. Predicted protein-coding genes are indicated with arrows, indicating the direction of transcription. ‘ami’ stands for N-actylmuramoyl-L-alanine amidase. CDS coding for transposases were annotated using ISFinder (Siguier, 2006). The length of the genomic island predicted by the IslandViewer package is indicated as well as the chromosomal origin of replication (ORI). Greyscale for similiarity levels by BLASTn is shown in the top right area of the figure.

**Figure 3.**
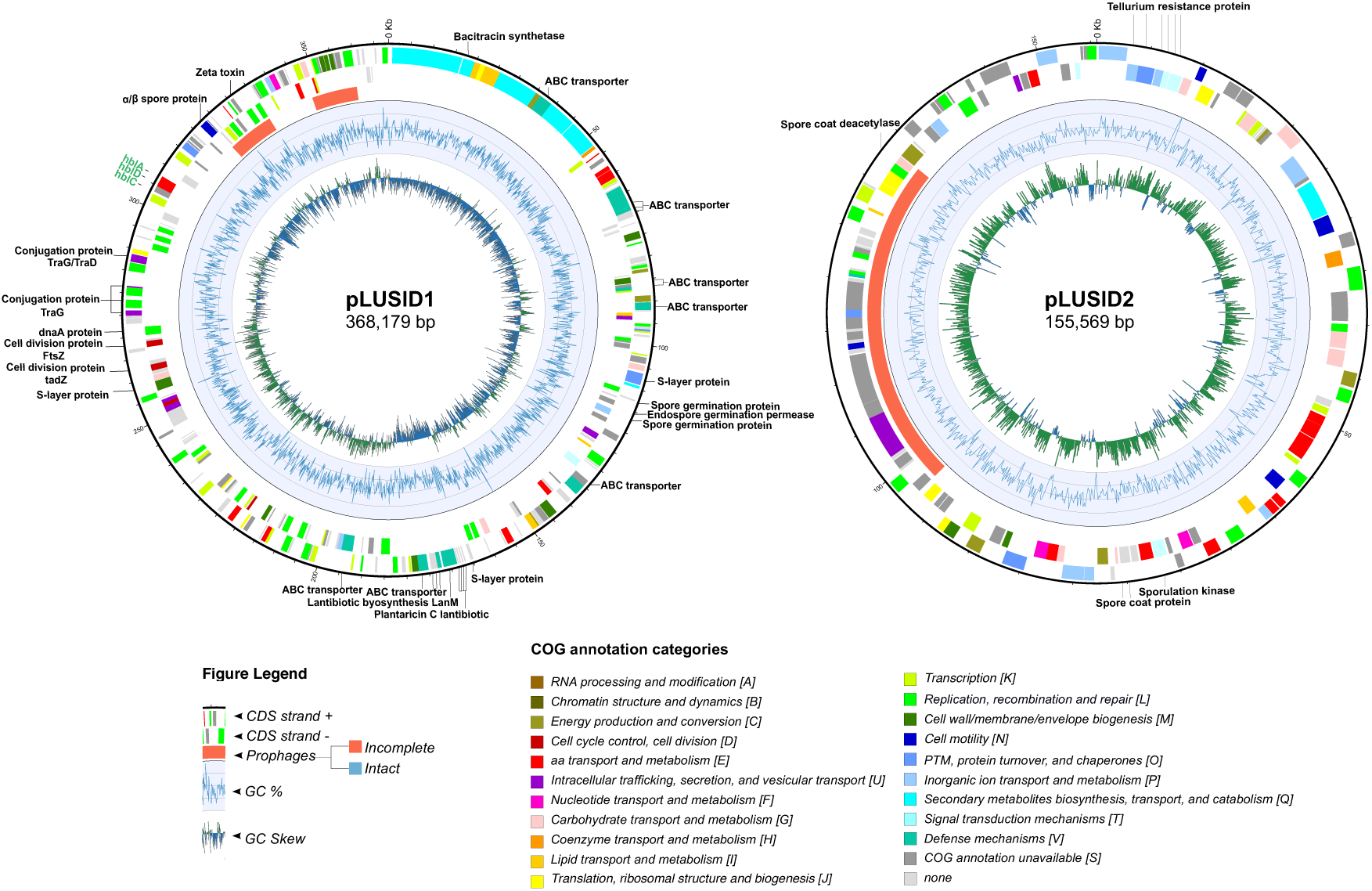
Circular representation of megaplasmids pLUSID1 and pLUSID2. The outer labels refer to toxin genes found using bTyper (green) and loci that are mentioned throughout the paper (black). The next two circles represent the coding regions on the positive and negative strands colored by their functional annotation. The next circle displays prophages predicted by PHASTER webserver where orange=incomplete, blue=intact. The inner two rings indicate the changes in %GC content and the GC skew respectively. Coordinates of the highlighted features can be found in Supplemental Table 3.

**Figure 4.**
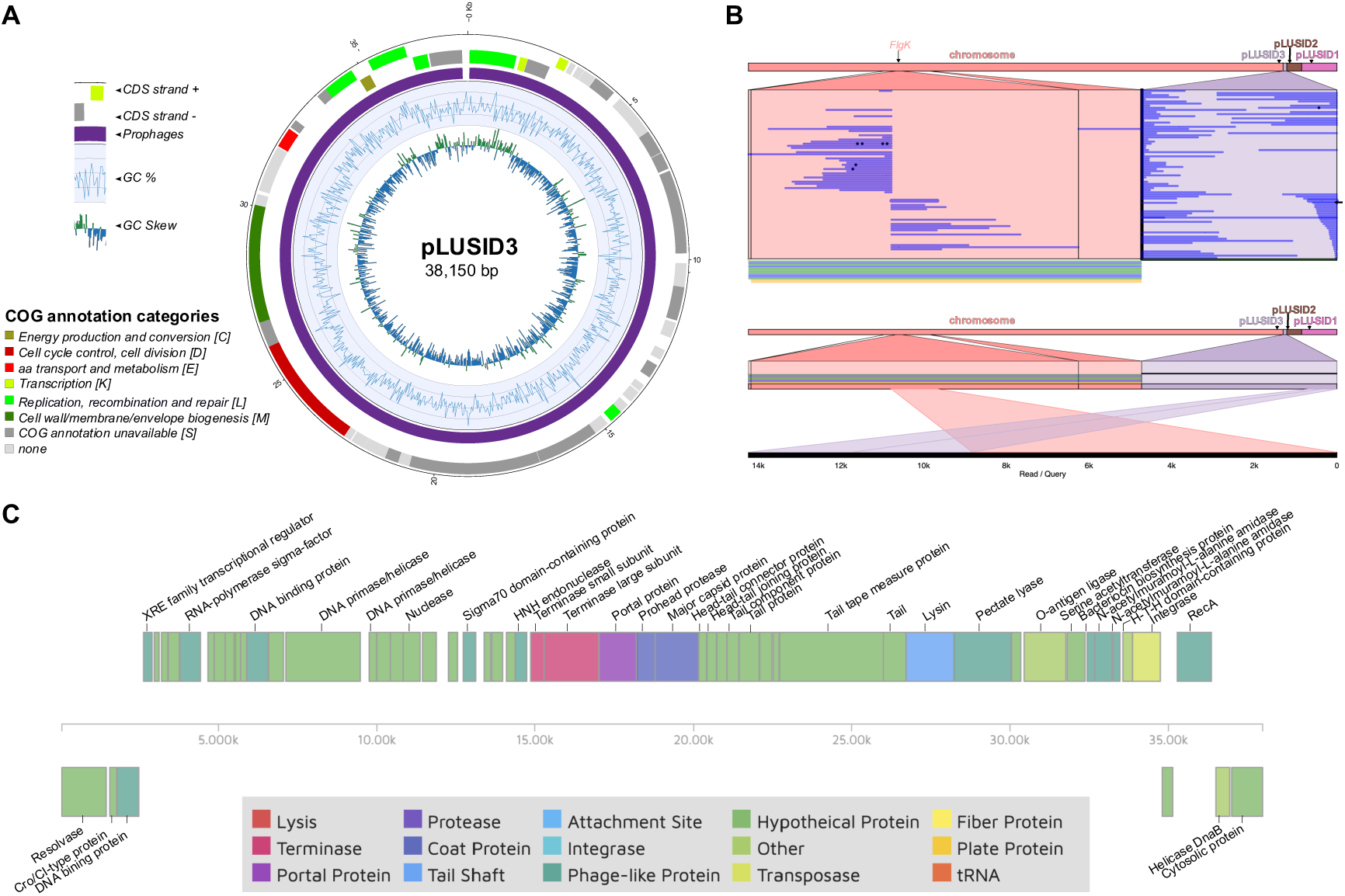
pLUSID3 features analysis (A) Circular representation of plasmid pLUSID3. The outer two circles represent the coding regions on the positive and negative strands colored by their functional annotation. The next circle displays prophages predicted by PHASTER webserver where purple=questionable. The inner two rings indicate the changes in %GC content and the GC skew respectively. (B) Structural variation analysis of pLUSID3. Long-reads from Nanopore sequencing show evidence for pLUSID3 variants in HER1410 genome where it can be integrated in the chromosome or form a closed circular element (upper panel). One of the reads showing the chromosomal integration is indicated by an arrow and represented in the lower panel. Variants are visualized with Ribbon (Nattestad et al., 2016). (C) CDS map provided and colored by the PHASTER tool. CDSs annotations were updated using PHASTER, PHMMER and COG functional annotations (Supplemental Table 4). Pale and dark green colors correspond to hypothetical and phage-related functions. Other colors indicate identified and annotated phage functions.

### LARGE STRUCTURAL VARIATION ANALYSIS

The larger structural variations were detected using in combination the long-reads mapper Ngmlr v0.2.7 and the structural variation caller Sniffles v1.0.11 (Sedlazeck et al., 2018). The results were visualized using the software Ribbon v1.1 (Nattestad et al., 2016).

### GENOME ANALYSIS

The identification of genomic islands of the annotated HER1410 genome was done using the Island Viewer 4 (Bertelli et al., 2017). The genomes were also mined for the presence of prophages using the web application PHASTER (Arndt et al., 2016). Comparative analysis of the *cry1Ba4*-containing genomic island with plasmidial *cry1Ba*-containing cassettes was performed and visualized using EasyFig (Sullivan et al., 2011).

The virulence and antibiotic resistance genes were annotated using BTyper version 2.3.3 (default settings) (Carroll et al., 2017) and insecticidal toxin-encoding genes were annotated using BTyper3 version 3.0.2 (Carroll et al., 2020). CRISPR-Cas genes were detected using CRISPRCasFinder (Couvin et al., 2018) (Supplemental Table 3). Chromosomal and extrachromosomal features were visualized using Circos (Krzywinski et al., 2009)

### PHYLOGENETIC ANALYSIS

To establish the position of *B. thuringiensis* HER1410 within the population structure of *B. cereus-thuringiensis*, two taxon sets were created. The first set (Supplemental Table 5) comprised up to five strains of *B. cereus* or *B. thuringiensis* selected from differentiated clades in BCSL_114 and *B. thuringiensis* phylogenetic trees from previous population structure analyses (Bazinet, 2017, Gillis et al., 2018). The second set comprised 1) the closest genomes to our strain identified using the phylogeny reconstruction of the first dataset (underlined in Figure 5), 2) 121 strains closely related to those according to phylogeny analysis of *B. cereus* group strains from Carroll et al. (2020), and 3) *B. thuringiensis* YBT-1518 type strain as outgroup (Supplemental Table 6). To ensure the assembly quality of both datasets, only genomes with an N50 size under 20 kbp were selected. The pangenomes of *B. cereus s. s.* and *B. thuringiensis* were inferred using Roary version 3.11.2 (Page et al., 2015). First, the GenBank files from selected strains were downloaded from NCBI’s and converted to GFF3 using the bp_genbank2gff script from the Bioperl library (Stajich et al., 2002). Second, the gff3 annotations were provided to Roary to calculate the pangenome of the dataset and produce a multiple sequence alignment (MSA) of the concatenated core genes (present in >99% strains) using MAFFT. Third, the MSA of the core genome was used to generate a best-fit maximum likelihood phylogeny using IQTREE version 1.6.12 using ModelFinder optimization (Nguyen et al., 2015, Kalyaanamoorthy et al., 2017). Finally, the trees were visualized in FigTree v.1.4.4 (Rambaut, 2012) and iTOL version 5.5 (Letunic and Bork, 2016).

**Figure 5.**
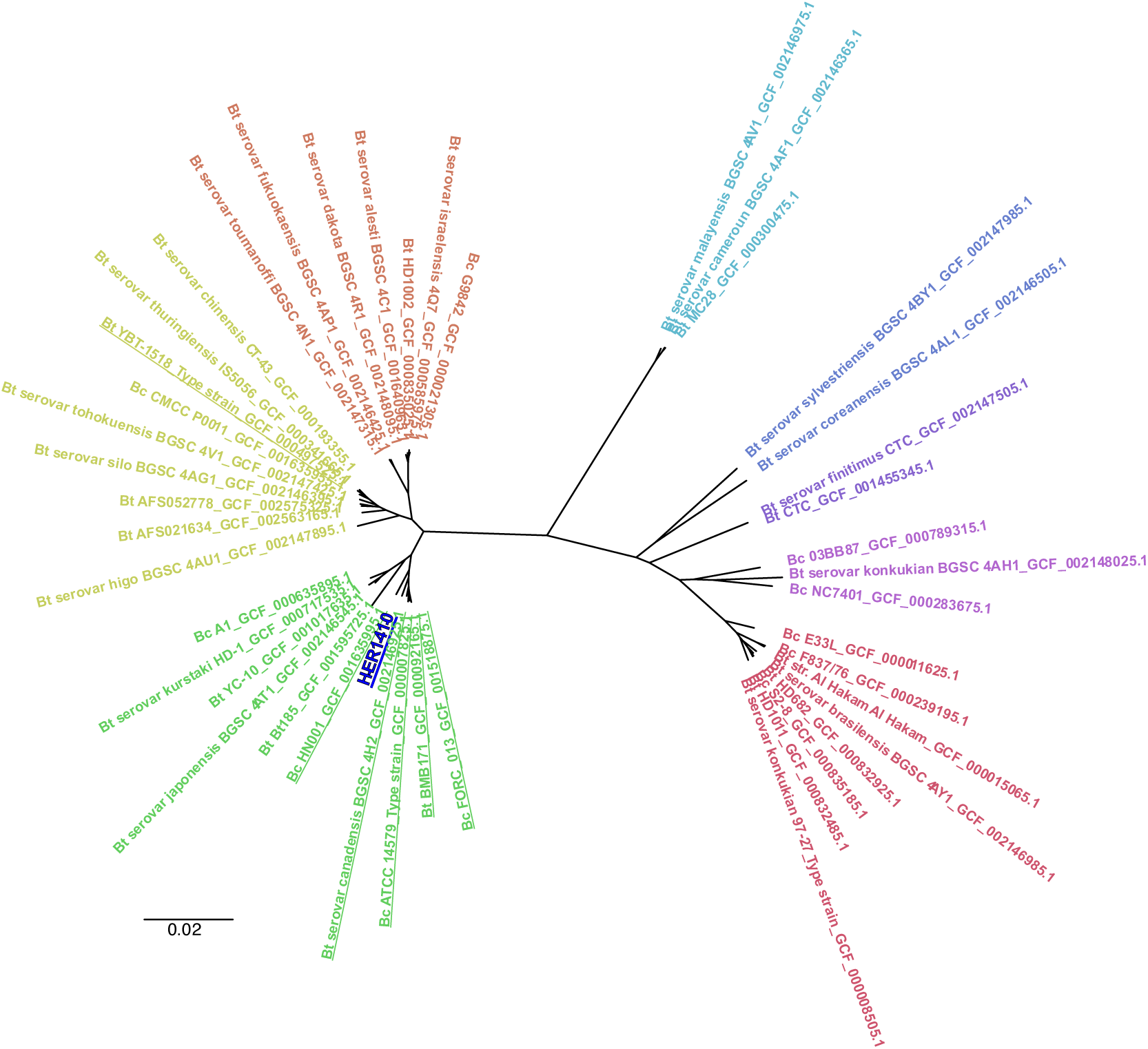
Overall position of *B. thuringiensis* HER1410 in the *Bacillus cereus-thuringiensis* phylogeny. An unrooted tree, representing the phylogenetic position of HER1410 (dark blue) among differentiated genomic sequences of the *B. thuringiensis* (Bt) and *B.cereus* (Bc) strains selected (Supplemental Table 5), was visualized with FigTree. A maximum likelihood tree was generated upon multiple sequence alignment of the core genome from selected strains. Labels correspond to their name in NCBI genomic database plus the assembly accession number. The colors represent clearly distinct clades. Strains in the same clade as HER1410 are underlined.

### DATA AVAILABILITY

The genome of *Bacillus thuringiensis* HER1410 was deposited under the NCBI GenBank accession numbers CP050183-CP050186S. The datasets of Illumina and Nanopore reads are available in the NCBI SRA database via the bioproject accession number PRJNA613019.

## Results and Discussion

### CHROMOSOME FEATURES

The *B. thuringiensis* HER1410 genome comprises a 5,585,577 bp circular chromosome which represents 90.86% of the whole genome. While gene content in chromosomes within the *B. cereus* group is well conserved, variation among them mainly stems from MGEs such as prophages, insertion elements, transposons and plasmids (Rasko et al., 2005). Accordingly, the search for genomic islands and prophages in HER1410 chromosome yielded 24 predicted genomic islands and nine prophages (three intact and six incomplete) that would belong to the *Caudovirales* (Figure 1, Supplemental Table 3). These prophages were assigned by Phaster software (Arndt et al., 2016) to the families *Siphoviridae* and *Myoviridae*, in agreement with previous studies of prophages infecting the *B. cereus* group (Gillis and Mahillon, 2014a). No tectiviruses were detected on the chromosome or the extrachromosomal elements, in line with the results from Verheust et al. (2005), which showed that no linear molecules could be detected by PFGE analysis of genomic DNA and where the sensitivity to tectiviral infections was confirmed.

*B. thuringiensis* is defined by its ability to produce crystal proteins with specific activity against insects. Crystal encoding genes in *B. thuringiensis* are harbored by plasmids (Ehling-Schulz et al., 2019). Only few chromosome-located exceptions have been reported in early works pre-dating the high-throughput sequencing era (Kronstad et al., 1983, Carlson and Kolsto, 1993). Strikingly, we found that HER1410 genome contains a single crystal protein-coding gene, *cry1ba4*, that is not located in a plasmid but within the chromosome, within a few kbp from the predicted origin of replication (Figure 2). This unusual location could be explained by the surrounding genetic context, a genomic island that contains multiple transposase genes from the IS*4* family. The association of transposases and *B. thuringiensis* toxins, in particular, transposases in the IS*4*, IS*6*, IS*66*, IS*605*, and Tn*3* families is extensively described and highlights the dynamics of horizontal transfer of toxins (Zheng et al., 2017). Within this island, several different transposases were annotated, suggesting multiple transposition events and pointing this region as a putative insertion hotspot (Figure 2). Interestingly, the HER1410 *cry1Ba4-*containing island organization resembles that of other *B. thuringiensis* plasmids where a N-acetylmuramoyl-l-alanine amidase CDS, coding for a germination key enzyme, precedes the insecticidal gene (Fiedoruk et al., 2017). When the HER1410 chromosomal *cry1Ba4-*containing island was compared with other plasmidial *cryBa1*-containing cassettes described in Fiedoruk et al. (2017), a similar genomic organization was observed (Figure 2). Also, as in *cry1Ba*-containing plasmids, the HER1410 island only posseses one pesticidal gene (not within a PAI) and it is not preceded by a K^þ^(Na^þ^)/H^þ^ antiporter. These similarities suggest a plasmidial origin of the HER1410 *cry1Ba4-*coding island. The insertion of this whole cassette into the chromosome close to the origin of replication may have been favored by a higher copy number of genes in this region during exponential growth, potentially allowing the loss of the plasmid (Bergstrom et al., 2000).

Since no additional crystal genes were detected, *cry1Ba4* can be related to the observation that a single type of parasporal crystal exists in this strain, a feature that allowed its classification as *B. thuringiensis* (Verheust et al., 2005). More than 700 genes encoding Cry proteins have been identified and characterized in *B. thuringiensis* Toxin Nomenclature (Crickmore, 2016) and they show highly specific insecticidal activity to target insects. In this case, Cry1Ba4 delta-toxin, purified from *B. thuringiensis* ser. *entomocidus* HD9, demonstrated mortality greater than 85% against *Plutella xylostella*, one of the more prevalent pests in Malaysia (Nathan et al., 2006). These results suggest that HER1410 may also possess insecticidal activity against some *Lepidoptera* insects.

Additional *B. cereus s. l.* toxins in this strain were found using the BTyper software. No emetic toxin cereulide, responsible of emetic syndrome, was detected, but heat-labile diarrheal syndrome enterotoxins cytolysin K (CytK), non-hemolytic enterotoxin (Nhe), and hemolysin BL (Hbl), commonly present among *B. cereus* group members, were found within the chromosome of HER1410 (Figure 1). Other virulence factors such as InhA2 and bceT are also present in the HER1410 genome. These results correlate with the hemolysis activity and cytotoxicity exhibited by this strain, opening up also the possibility that HER1410 might cause diarrheal syndrome. However, it is difficult to predict the enterotoxic potential of a given strain from genomics data only since it stems from a complex mechanism where transcription levels and other virulence factors are involved (Pruss et al., 1999a, Ehling-Schulz et al., 2019). The presence of antibiotic resistance genes and CRISPR-Cas systems was also searched, using BTyper and CRISPRCasFinder tools. Several vancomycin, beta-lactam and fosfomicin resistance genes but no CRISPR systems were detected along the chromosome (Figure 1, Supplemental Table 3).

### THREE NOVEL EXTRACHROMOSOMAL ELEMENTS IN *BACILLUS THURINGIENSIS*

Extrachromosomal elements in *B. cereus s. l.* are numerous, variable, mobile, and essential to the ecology and classification of its members. In particular, some plasmids are extensively studied since they carry virulence genes like anthrax containing pXO1 (182 kb) and pXO2 (95 kb) in *B. anthracis* or crystal toxins containing megaplasmids characteristic of *B. thuringiensis* (Adams et al., 2014). Despite the importance of plasmids in this group, the common use of short-reads sequencing techniques has often resulted in draft genome assemblies lacking described extrachromosomal elements (Meric et al., 2018). In this case, nanopore long-reads sequencing provided an efficient solution to identify two megaplasmids, denominated pLUSID1 (368 kb), pLUSID2 (155 kb), and a smaller plasmid, pLUSID3 (38 kb), comprising 9.14% of the genomic information. We used these three plasmids as queries in nucleotide-nucleotide BLASTn against the database of nucleotide collection (nr/nt) on NCBI. Based on BLASTn, pLUSID1 could be linked to some *B. thuringiensis* and *B. cereus s. s.* plasmids, being the closer homolog *B. thuringiensis* strain YC-10 plasmid pYC1 (70% coverage and 98.78% identity; Genbank CP011350.1). On the other hand, pLUSID2 and pLUSID3 hits corresponded mainly with *B. thuringiensis* and *B. cereus s. s.* chromosomes and, in the case of pLUSID3, BLASTn also highlighted a similarity to some *Bacillus* phages (see below).

The identification of pLUSID1, a megaplasmid of 368,179 bp, is in agreement with the previous report of direct observation by gel electrophoresis of a megaplasmid in HER1410 in Gillis et al. (2016). This element presents a lower GC-content than the chromosome and is detected in a low copy number of 1.0 ± 0.14 per chromosome, as is generally observed for large plasmids (Supplemental Table 1) (Bolotin et al., 2017). Within the *B. cereus* group, genes on plasmids are generally more similar to the chromosomal variable genes than to the chromosomal core genes. Also, these plasmids show some differences from chromosomes in the functions of the genes they harbor (Zheng et al., 2015). Comparing with the chromosome, the pLUSID1 functional annotation revealed an enrichment in defense mechanism genes and in secondary metabolite biosynthesis, transport and catabolism genes which are located in a particular region of the plasmid (Figure 3; Supplemental Figure 2). Besides, 20.54% of the annotated genes are related to replication, recombination, and repair representing the largest proportion of plasmid genes. Among them, several transposases and integrases were identified along with the plasmid. Two genomic islands were identified, one of them coinciding with an incomplete prophage. Also, within this category, a region located in one of the GC-skew changes contains some putative DnaA and cell division proteins (Ftsz-like protein and TadZ) (Figure 3, Supplemental Table 3). Similarly to the toxin-encoding plasmid *B. thuringiensis* pBtoxis, this region may contain the replication origin of pLUSID1 and these proteins could be responsible for a self-replication system and possibly the segregation and partitioning of this plasmid to daughter cells (Tang et al., 2007). Alternatively, this region might also contain a conjugation origin, as several conjugal transfer proteins were annotated in this location. Moreover, we could predict an S-layer protein, involved in plasmid mobilization (Gillis et al., 2018). Overall, these results suggest that pLUSID1 could be transferred by conjugation.

Unlike other *B. thuringiensis* megaplasmids, this element does not contain insecticidal toxin genes, but it harbors the three Hbl enterotoxin components, which means an additional copy to the chromosomal Hbl. Some antimicrobial toxins are also located in this plasmid. Thus, CDS coding for a bacitracin, a zeta toxin and four copies of a plantaricin C family lantibiotic as well as several ABC transporters, reportedly involved in bacteriocin secretion, were found (Berry et al., 2002). pLUSID1 also encodes spore-related proteins that contribute to the germination process. Homologous proteins have been found in *Clostridium botulinum* where they act as germination receptors that have been shown to be dispensable (Clauwers et al., 2017). Well studied toxin-encoding plasmids such as *B. anthracis* pXO1 and pBtoxis also encode genes involved in sporulation and germination and they have shown to be important for the virulence of the host (Guidi-Rontani et al., 1999, Berry et al., 2002). Overall, although pLUSID1 does not carry pesticidal proteins, it seems to have a role in HER1410 virulence and defense.

A second megaplasmid, pLUSID2 (155,569 bp), was found in HER1410 (Figure 3). This plasmid has not been experimentally observed for this strain and BLASTn analysis yielded high similarity to other strains’ chromosomes, where the closer homologue corresponded to the *B. thuringiensis* serovar *tolworthi* chromosome (94% coverage and 99.09% identity; Genbank AP014864.1). Nevertheless, as a result of the structural variation analysis of pLUSID2, no variants that could indicate this region is part of the chromosome, could be detected. Furthermore, the only structural variant observed is a result from the linearization of this plasmid (Supplemental Figure 3). Thus, the nanopore data supports that pLUSID2 is a circular element present 2 .0 ± 0.37 times per chromosome (Supplemental Table 1). This could mean that pLUSID2 is a cryptic plasmid in HER1410 and its high similarity to other *B. thuringiensis* chromosomes is consistent with the high mobility of MGE between plasmids and chromosomes within the *B. cereus* group (Meric et al., 2018). In agreement with that, several transposases integrases and recombinases were detected. However, this plasmid contains only one mobile island that coincides with the location of an incomplete prophage (Figure 3, Supplemental Table 3). BTyper results did not yield any *B.cereus*-related toxins or antibiotic resistance genes suggesting pLUSID2 is not directly associated with HER1410 virulence. Thus, pLUSID2 seems to play some role in sporulation, since it does possess four sporulation genes as well as six tellurite resistance proteins which have been shown in *B. cereus s. l.* spores (Delvecchio et al., 2006). Gene functional analysis also suggests a possible role related to basal metabolism, such as those involved in carbohydrate transport and metabolism, and inorganic ion transport and metabolism and energy production and conversion whose proportion seems to be higher than on the chromosome (Supplemental Figure 2, Supplemental Table 2). This contrasts with the general characteristics of *B. cereus* plasmids where the proportion of gene families involved in basal metabolism was significantly lower than on chromosomes (Zheng et al., 2015).

The smallest extrachromosomal element, pLUSID3, is a plasmid of 38,150 bp and coverage of 2.33x compared to the chromosome (Figure 4A, Supplemental Table 1). Annotation of pLUSID3 resulted in 40% of hypothetical proteins and a high number of phage related proteins (Supplemental Table 2). This annotation was updated with results from COG functional analysis (Huerta-Cepas et al., 2017), PHASTER (Arndt et al., 2016) and PHMMER tools (Potter et al., 2018) (Figure 4C, Supplemental Table 4). PHASTER results identified nearly the entire length of the plasmid as a questionable prophage, related to *Bacillus* prophage phBC6A52 (Genbank NC_004821.1). However, only 27% of pLUSID3 displays similarity to this prophage. The whole sequence of pLUSID3, as well as the annotated terminase and major capsid protein, were used as queries for BLASTn and BLASTp respectively against tailed phages NCBI database (taxid:28883). Results yielded similarity of pLUSID3 with phage Wrath (40% coverage, 83.63% identity) a *Bacillus* siphovirus found in the human bladder (Garretto et al., 2019). These results suggest that pLUSID3 is a circular plasmidial prophage, not previously reported, showing partial similarity to characterized *B. cereus* prophages. This is not surprising given the high diversity exhibited by prophages in *B. thuringiensis* (Fu et al., 2019). Strikingly, the structural variation analysis based on the nanopore data allowed us to show that pLUSID3 has a dual lifestyle and can be fully integrated into the chromosome and also persist as an extrachromosomal element (Figure 4B). Functional annotation revealed the presence of three different recombinases located in the same region of the plasmid. These recombinases may participate in the integration of this putative prophage into the chromosome, and may serve to resolve replication intermediates. It is remarkable that, in the limited clonal community of bacteria used for sequencing, we could detect several integration events of the whole plasmidial prophage that suggest a high mobility rate of this element. Interestingly, the integration was always detected in the same region of the chromosome, corresponding to the location of a flagellar hook associated protein CDS (*FlgK*). It has already been shown that the induction of the integration/excision of prophages in *B. thuringiensis* can modify some bacterial features. This is the case, for example, of phIS3501 that disrupts the production of hlyII toxin when it integrates into the chromosome (Moumen et al., 2012). FlgK is an essential element of the hook-filament junction and its absence causes flagellar misassembly and accumulation in the extracellular milieu of *Bacillus subtilis* (Cairns et al., 2014). The intermittent disruption of this gene by pLUSID3 integration could have an impact on the motility of *B. thuringiensis* HER1410 and biofilm formation (Houry et al., 2010). Together with the presence of putative cro/cI type transcriptional repressors, these results point to a hypothetical mechanism superinfection exclusion. Also, this set of cro/cI type transcriptional regulator genes grouped in the same region could be responsible for the SOS response mediated induction of this lysogenic phage.

### PHYLOGENETIC POSITIONING OF HER1410

The lack of HER1410 genomic information has resulted in an unsteady and uncertain classification of this strain. HER1410 was first classified as *B. thuringiensis* ser. *israelensis*, but was repositioned within the serovar *thuringiensis* (Verheust et al., 2005). Several studies have proposed to resolve the conflict between genomics and phenotypical classification (Bazinet, 2017, Baek et al., 2019, Carroll et al., 2020). While *B. thuringiensis* has always been characterized by their ability to produce entomopathogenic crystals, some crystal producing strains are more genetically similar to *B. cereus s. s.* (Carroll et al., 2020).

Here we aimed to characterize HER1410, not only by its capacity of producing entomotoxins, but also by its genomic features. Multiple different approaches have been used to stablish the population structure of *B. cereus s. l*. such as multi-locus sequence typing, ANI or core genes-based phylogenies (Bolotin et al., 2017, Baek et al., 2019, Carroll et al., 2020). Here, in order to provide a high resolution on the population structure, we performed a phylogenomics analysis based on a MSA of the core genes present in the pangenome (Page et al., 2015). To accomplish this using a comprehensive dataset, we applied a two-stage phylogenetic analysis. First, we selected 45 genomically differentiated *B. cereus s. s.* and *B. thuringiensis* strains to situate this HER1410 among defined clusters of *B. cereus* (see Methods). The pangenome of the selected *B. cereus-thuringiensis* strains was determined with Roary and resulted in a total of 1,838 core genes (i.e. present in 99% of the strains), which were subsequently aligned to build a ML phylogenetic tree using IQ-Tree (Figure 5). We could observe that there are some clusters including both *B. cereus* and *thuringiensis*, in agreement with the reported high genomic similarities between both species. Interestingly, although HER1410 has been classified as *B. thuringiensis* ser. *thuringiensis*, our results locate this strain in a cluster that comprises both *B. cereus s. s.* and *thuringiensis* strains. Among these strains, we could also find the type strain for *B. cereus s. s.*, ATCC 14579.

Furthermore, to find the closest genomes to our strain, we performed a focused analysis using the obtained similar strains to HER1410. These strains belong to the same cluster within the phylogeny analysis of *B. cereus* group from a recent work on its population structure (Carroll et al., 2020). Thus, we selected the 126 strains included in this cluster, with the addition of HER1410, and *B. thuringiensis* YBT-1518 type strain as outgroup and obtained the pangenome of this dataset, with a total of 2,886 core genes. This deeper analysis accurately anchored HER1410 within a clade which contains both *B. cereus* and *thuringiensis* strains (Figure 6). These results agree with the taxonomic classification using BTyper3 of HER1410 as *B. cereus s. s.* biovar *thuringiensis*. This software attempts to reconcile genomic definitions of species with clinically and industrially relevant phenotypes (Carroll et al., 2020). In this case, HER1410 could be classified as *Bacillus cereus s. s.* because of its similarity to this genomospecies (ANI=98.08%) and biovar *thuringiensis* due to the presence of the insecticidal gene *cry1Ba4*. For practical purposes and, in line with the currently accepted nomenclature, we propose maintaining HER1410 as a *B. thuringiensis* serovar *thuringiensis* strain although the separation between these two species does not fully hold upon genomic inspection.

**Figure 6.**
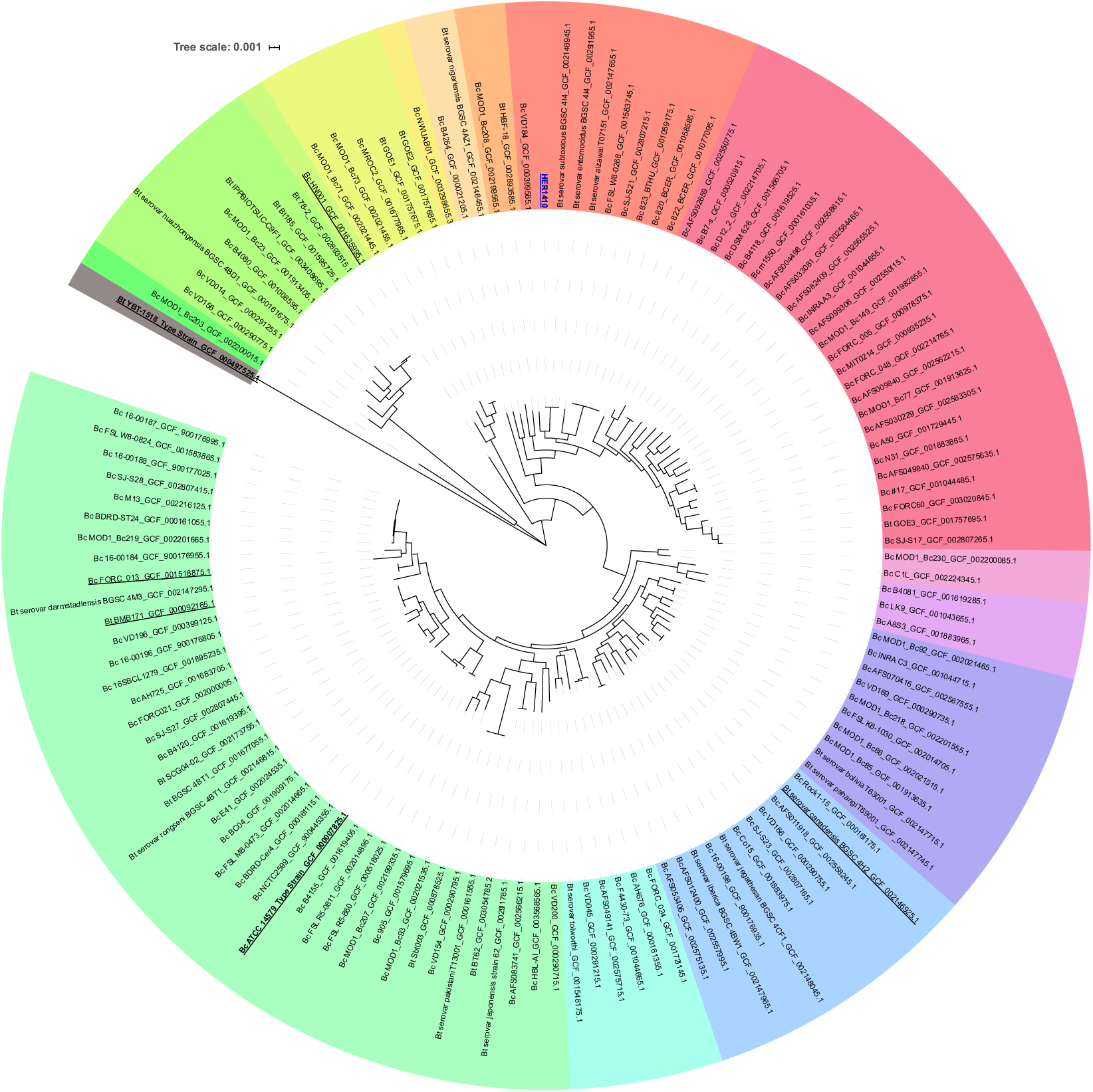
Accurate positioning of *B. thuringiensis* HER1410 in the *Bacillus cereus-thuringiensis* phylogeny. Maximum likelihood rooted tree representing the phylogeny of differentiated genomic sequences of the *B. thuringiensis* (Bt) and *B.cereus* (Bc) strains selected (Supplemental Tables 6) was visualized with iTOL. *Bt* YBT-1518 was used as an outgroup for pangenome analysis and core genome alignment and, subsequentially, to root the obtained tree. Labels correspond to their name in NCBI genomic database plus the assembly accession number. The colors represent clearly distinct clades. Common strains in the two phylogenetical analysis (Figures 5 and 6) are underlined and the strain of interest, HER1410 is colored in blue.

Interestingly, *B. thuringiensis* HER1410 has a high level of similarity with *B. thuringiensis* ser. *subtoxicus* and ser. *entomocidus*. The latter has shown an antimicrobial activity against Gram-positive bacteria including *Listeria monocytogenes*, one of four pathogenic *Pseudomonas aeruginosa* and several fungi. Despite the presence of a range of virulence-related genes, including Hbl, ser. *entomocidus* is not toxic against mammalian Vero cells (Cherif et al., 2003). This would not be the case for HER1410 which does present cytotoxic activity. Also, the *entomocidus* genome contains more than one *cry* gene besides some other entomocidal toxins not present in HER1410 suggesting both strains, although very similar, show different virulence phenotypes (Meric et al., 2018). Interestingly, as reported here for HER1410, *entomocidus* has been suggested to harbor *cry* genes within the chromosome (Carlson and Kolsto, 1993). On the other hand, *cry* genes also seemed to be present in plasmids for *entomocidus.* This could explain the different number of *cry* genes for each strain by the presence of a *cry*-encoding plasmid in *entomocidus* but not in HER1410. Because the two closer strains (*entomocidus* and *subtoxicus*) have been sequenced using Illumina, the quality of the resulting assemblies did not allow the identification of extrachromosomal elements. However, BLASTn analysis indicate pLUSID1, pLUSID2 and pLUSID3 are also present in these strains.

As mentioned above, besides HER1410, two other strains, HER1047 and GBJ002, are broadly used to the study of tectiviruses because of their sensitivity to multiple *Betatectivirus* members. This is in conflict with the hypothesis that these phages are probably able to infect only specific bacterial strains from closely related hosts, unlike the PRD1-like phages that have a broad host range (Gillis and Mahillon, 2014b). On one hand, HER1047 is classified as *B. cereus*, as it does not produce crystals but contains the *B. cereus* gene *clo* (Verheust et al., 2005) while, on the other hand, GBJ002 is derived from *B. thuringiensis* ser. *israelensis* 4Q7, which is cured of its plasmids. Interestingly, HER1410 and *israelensis* 4Q7 seems to be a distant relative in our phylogenetical analysis. This is consistent with results obtained in Gillis and Mahillon (2014b), where it was not possible to pinpoint a clear association between the phage infection pattern and the *Bacillus* species. The sequencing of both GBJ002 and HER1047 would allow a comparative analysis to obtain some clues about their common features that could establish a pattern for tectiviral susceptibility.

### CONCLUDING REMARKS

Using a combination of short and long-reads sequencing methods has proven to be particularly useful to unravel important features in a bacterial species where delineation of plasmids and location of genes is key. Our high-quality assembly of *B. thuringiensis* HER1410 genome led to the identification of an uncommon location of a *cry* gene, close to the replication origin of the chromosome rather than within a plasmid, highlighting the high mobility of these genes. The implications of this unusual location should be studied further as it may have an effect on the entomopathogenicity of this strain against lepidoptera insects and, therefore, could make HER1410 a promising candidate as a specific biopesticide.

Resolving the genome has also allowed the identification of three plasmids not reported previously. In particular, the identification of a small integrative plasmidial prophage related to a *Bacillus* virus found in the human bladder and its role in flagellar gene disruption has a great interest in the *Bacillus* phages research field and their interaction with the human microbiome.

Finally, the genomic characterization of HER1410 will be valuable for studies on *Bacillus* phages. A better understanding of this highly sensitive strain to phages, especially to tectiviruses, may improve further investigations on virus-host interactions and *Bacillus* phages characterization.

## Acknowledgments

This research was funded by Fundación Ramón Areces (XIX Concurso Nacional de Investigación en Ciencias de la Vida y la Materia) and institutional grants from Fundación Ramón Areces and Banco de Santander to the Centro de Biología Molecular Severo Ochoa. A. L. is holder of a PhD fellowship [FPU15/05797] from the Spanish Ministry of Science, Innovation and Universities.

C. L. is supported by a PhD fellowship from FWO Vlaanderen (1S64720N).

The authors would also like to thank Dr. Annika Gillis for her critical input on the content of the manuscript.

## Supplemental data

**Supplemental Figure 1.**
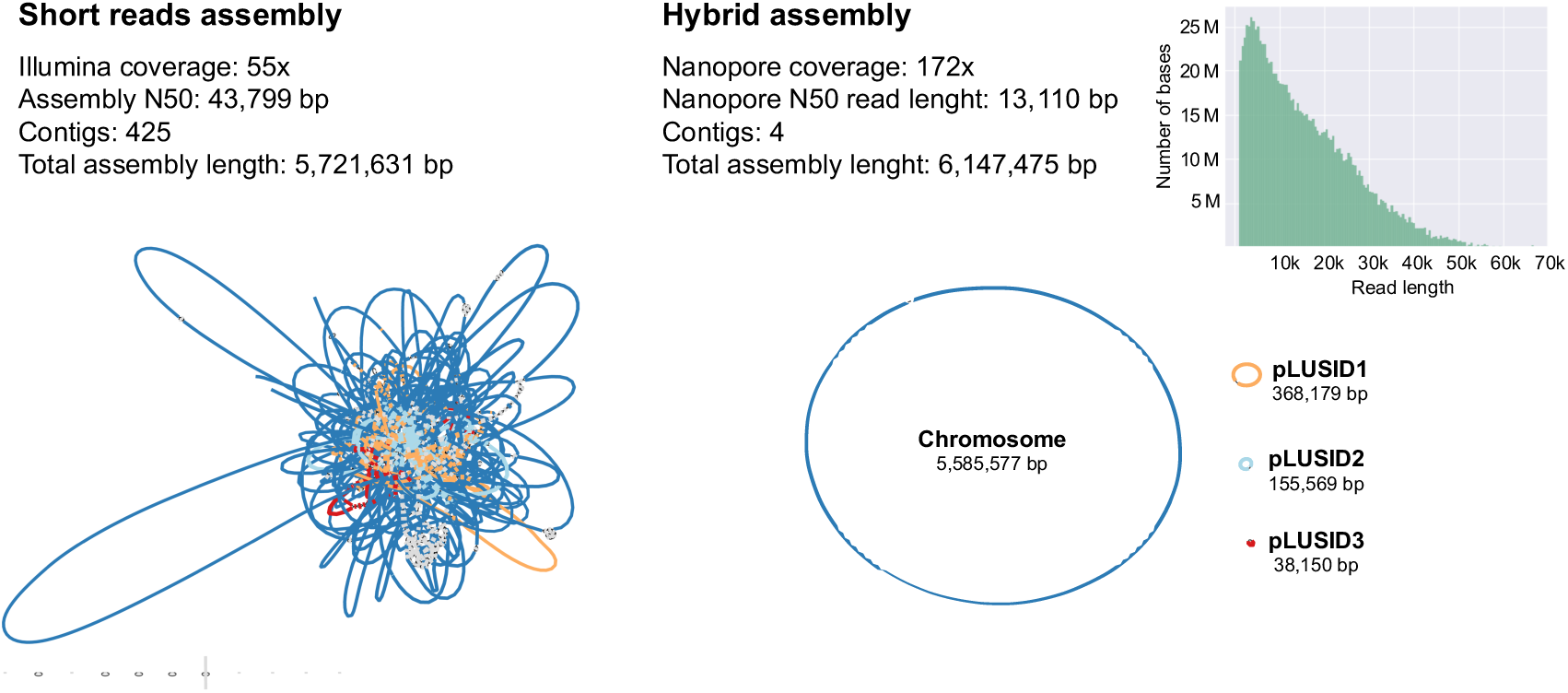
Comparative metrics of short reads assembly using Illumina sequences and hybrid assembly using Illumina and Nanopore sequences. Representation of assemblies was created with Bandage where chromosome=dark blue, pLUSID1=orange, pLUSID2=light blue, pLUSID3=red.

**Supplemental Figure 2.**
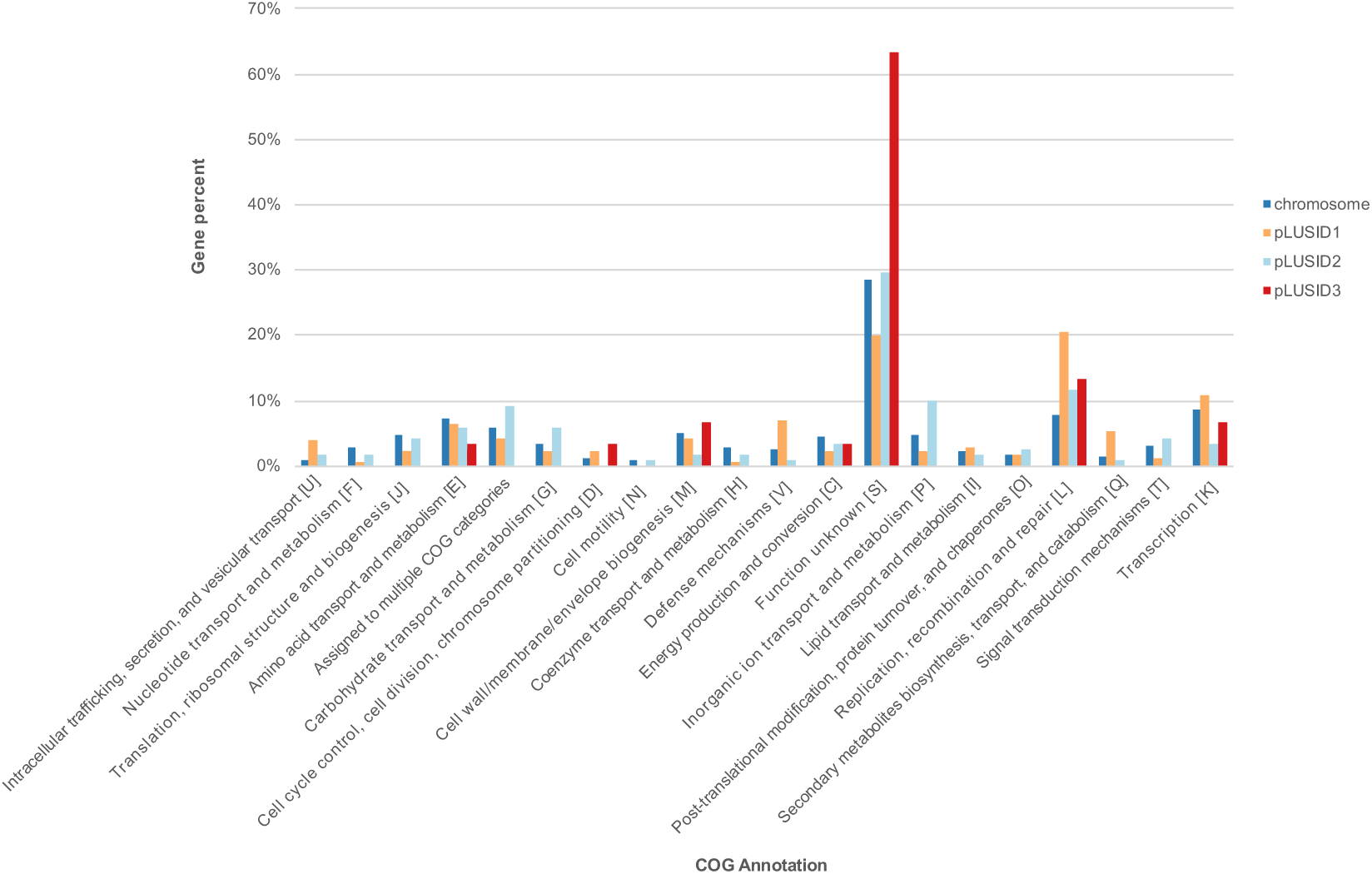
Bar plot of COG categories proportions for the elements of HER1410 genome. Ratio of genes belonging to each COG category, relative to the total number of genes in each element is shown (Supplemental Table 2).

**Supplemental Figure 3.**
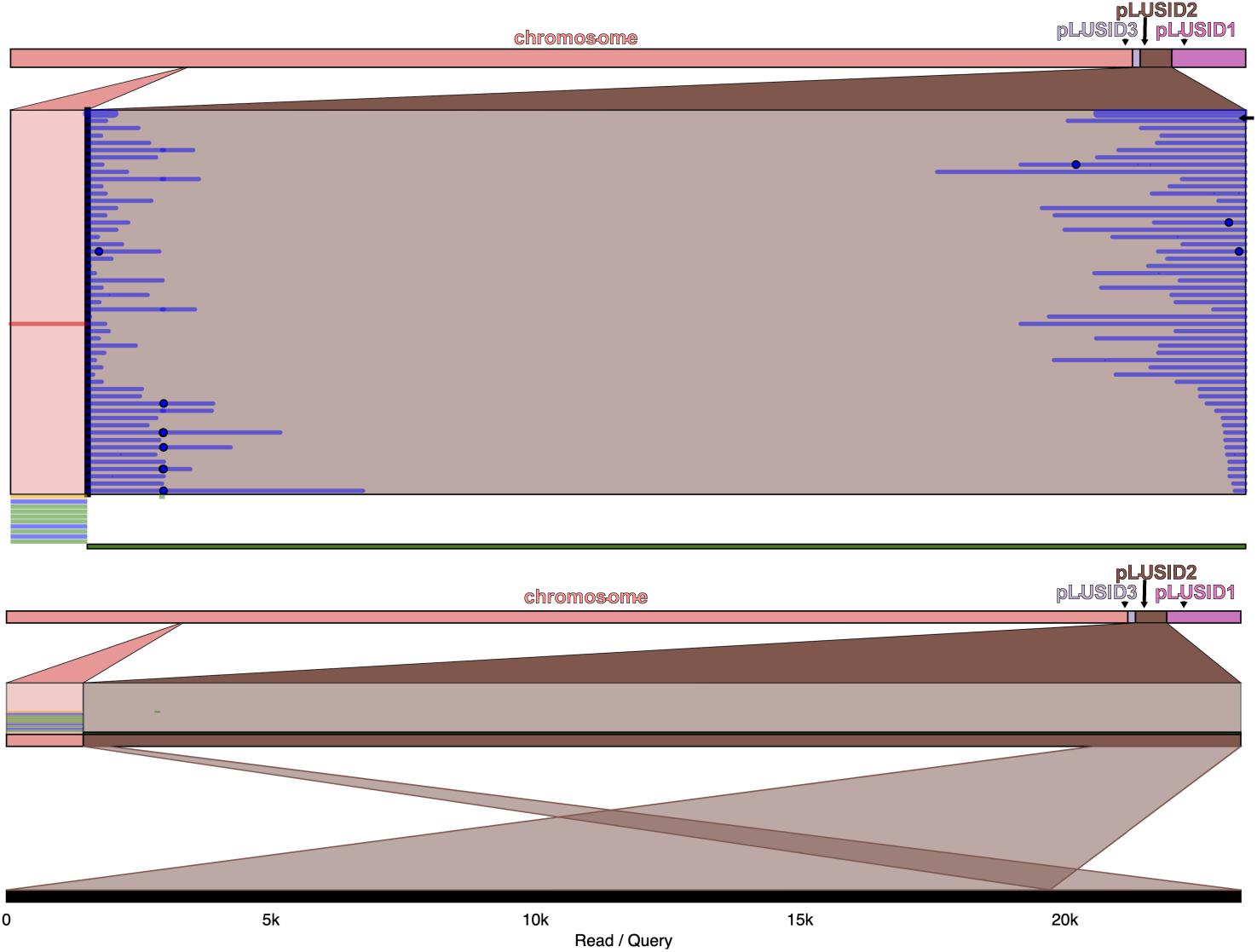
Structural variation analysis of pLUSID2. Long reads from Nanopore sequencing show evidence for pLUSID2 in HER1410 genome as a closed circular element. The only structural variation detected shown above resulted from the linearization of the circular molecule sequence (upper panel). One of the reads is indicated by an arrow and represented in the lower panel. Variants are visualized with Ribbon (Nattestad et al., 2016).

**Table S1.** HER1410 genome assembly statistics.

**Table S2.** Annotation information and number and percentage of genes assigned to COG categories for HER1410 genome.

**Table S3.** Chromosomal and extrachromosomal features detected in this work (Figures 1, 2 and 3).

**Table S4.** Detailed annotation of pLUSID3. Closest homolog for PGAP, eggNOGmapper, PHMMER and PHASTER are shown for each locus.

**Table S5.** Strains dataset used for *Bacillus cereus sensu lato* unrooted phylogenetic tree (Figure 5).

**Table S6.** Strains dataset used for HER1410-related strains phylogenetic tree (Figure 6).

## References

Adams, V., Li, J., Wisniewski, J. A., Uzal, F. A., Moore, R. J., et al., 2014. Virulence plasmids of spore-forming bacteria. Microbiol. Spectrum. 2(6): PLAS-0024-2014. https://doi.org/10.1128/microbiolspec.plas-0024-2014

Albers, P., Lood, C., Ozturk, B., Horemans, B., Lavigne, R., et al., 2018. Catabolic task division between two near-isogenic subpopulations co-existing in a herbicide-degrading bacterial consortium: Consequences for the interspecies consortium metabolic model. Environ. Microbiol. 20(1): 85–96. https://doi.org/10.1111/1462-2920.13994

Andrews, S., 2010. Fastqc: A quality control tool for high throughput sequence data. Available: https://www.bioinformatics.babraham.ac.uk/projects/fastqc/.

Arndt, D., Grant, J. R., Marcu, A., Sajed, T., Pon, A., et al., 2016. Phaster: A better, faster version of the phast phage search tool. Nucleic Acids Res. 44(W1): W16–21. https://doi.org/10.1093/nar/gkw387

Baek, I., Lee, K., Goodfellow, M. & Chun, J., 2019. Comparative genomic and phylogenomic analyses clarify relationships within and between *Bacillus cereus* and *Bacillus thuringiensis*: Proposal for the recognition of two *Bacillus thuringiensis* genomovars. Front. Microbiol. 10:1978. https://doi.org/10.3389/fmicb.2019.01978

Bankevich, A., Nurk, S., Antipov, D., Gurevich, A. A., Dvorkin, M., et al., 2012. Spades: A new genome assembly algorithm and its applications to single-cell sequencing. J. Comput. Biol. 19(5): 455–477. https://doi.org/10.1089/cmb.2012.0021

Bazinet, A. L., 2017. Pan-genome and phylogeny of *Bacillus cereus* sensu lato. BMC Evol. Biol. 17(1): 176. https://doi.org/10.1186/s12862-017-1020-1

Bergstrom, C. T., Lipsitch, M. & Levin, B. R., 2000. Natural selection, infectious transfer and the existence conditions for bacterial plasmids. Genetics 155(4): 1505–19.

Berjon-Otero, M., Villar, L., Salas, M. & Redrejo-Rodríguez, M., 2016. Disclosing early steps of protein-primed genome replication of the gram-positive tectivirus Bam35. Nucleic Acids Res. 44(20): 9733–9744. https://doi.org/10.1093/nar/gkw673

Berry, C., O’Neil, S., Ben-Dov, E., Jones, A. F., Murphy, L., et al., 2002. Complete sequence and organization of pbtoxis, the toxin-coding plasmid of *Bacillus thuringiensis* subsp. *israelensis*. Appl. Environ. Microbiol. 68(10): 5082–5095. https://doi.org/10.1128/aem.68.10.5082-5095.2002

Bertelli, C., Laird, M. R., Williams, K. P., Simon Fraser University Research Computing, G., Lau, B. Y., et al., 2017. Islandviewer 4: Expanded prediction of genomic islands for larger-scale datasets. Nucleic Acids Res. 45(W1): W30–W35. https://doi.org/10.1093/nar/gkx343

Bolger, A. M., Lohse, M. & Usadel, B., 2014. Trimmomatic: A flexible trimmer for illumina sequence data. Bioinformatics 30(15): 2114–20. https://doi.org/10.1093/bioinformatics/btu170

Bolotin, A., Gillis, A., Sanchis, V., Nielsen-LeRoux, C., Mahillon, J., et al., 2017. Comparative genomics of extrachromosomal elements in *Bacillus thuringiensis* subsp. israelensis. Res Microbiol. 168(4): 331–344. https://doi.org/10.1016/j.resmic.2016.10.008

Botelho, J., Lood, C., Partridge, S. R., van Noort, V., Lavigne, R., et al., 2019. Combining sequencing approaches to fully resolve a carbapenemase-encoding megaplasmid in a pseudomonas shirazica clinical strain. Emerg Microbes Infect. 8(1): 1186–1194. https://doi.org/10.1080/22221751.2019.1648182

Cairns, L. S., Marlow, V. L., Kiley, T. B., Birchall, C., Ostrowski, A., et al., 2014. Flgn is required for flagellum-based motility by *Bacillus subtilis*. J. Bacteriol. 196(12): 2216–2226. https://doi.org/10.1128/jb.01599-14

Carlson, C. R. & Kolsto, A. B., 1993. A complete physical map of a *Bacillus thuringiensis* chromosome. J. Bacteriol. 175(4): 1053–60. https://doi.org/10.1128/jb.175.4.1053-1060.1993

Carroll, L. M., Kovac, J., Miller, R. A. & Wiedmann, M., 2017. Rapid, high-throughput identification of anthrax-causing and emetic *Bacillus cereus* group genome assemblies via btyper, a computational tool for virulence-based classification of *Bacillus cereus* group isolates by using nucleotide sequencing data. Appl. Environ. Microbiol. 83(17): e01096–17. https://doi.org/10.1128/aem.01096-17

Carroll, L. M., Wiedmann, M. & Kovac, J., 2020. Proposal of a taxonomic nomenclature for the *Bacillus cereus* group which reconciles genomic definitions of bacterial species with clinical and industrial phenotypes. mBio. 11(1): e00034–20. https://doi.org/10.1128/mbio.00034-20

Clauwers, C., Lood, C., Van den Bergh, B., van Noort, V. & Michiels, C. W., 2017. Canonical germinant receptor is dispensable for spore germination in *Clostridium botulinum* group ii strain nctc 11219. Sci. Rep. 7(1): 15426. https://doi.org/10.1038/s41598-017-15839-y

Couvin, D., Bernheim, A., Toffano-Nioche, C., Touchon, M., Michalik, J., et al., 2018. Crisprcasfinder, an update of crisrfinder, includes a portable version, enhanced performance and integrates search for cas proteins. Nucleic Acids Res. 46(W1): W246–W251. https://doi.org/10.1093/nar/gky425

Crickmore, N., Feitelson, J., 2016. Bacillus thuringiensis toxin nomenclature. Available: http://www.lifesci.sussex.ac.uk/Home/Neil_Crickmore/Bt/

Daugelavicius, R., Gaidelyte, A., Cvirkaite-Krupovic, V. & Bamford, D. H., 2007. On-line monitoring of changes in host cell physiology during the one-step growth cycle of Bacillus phage Bam35. J. Microbiol. Methods, 69(1): 174–9. https://doi.org/10.1016/j.mimet.2006.12.023

De Coster, W., D’Hert, S., Schultz, D. T., Cruts, M. & Van Broeckhoven, C., 2018. Nanopack: Visualizing and processing long-read sequencing data. Bioinformatics 34(15): 2666–2669. https://doi.org/10.1093/bioinformatics/bty149

Delvecchio, V. G., Connolly, J. P., Alefantis, T. G., Walz, A., Quan, M. A., et al., 2006. Proteomic profiling and identification of immunodominant spore antigens of *Bacillus anthracis, Bacillus cereus*, and *Bacillus thuringiensis*. Appl. Environ. Microbiol. 72(9): 6355–6363. https://doi.org/10.1128/aem.00455-06

Ehling-Schulz, M., Lereclus, D. & Koehler, T. M., 2019. The *Bacillus cereus* group: *Bacillus* species with pathogenic potential. Microbiol. Spectr. 7(3). https://doi.org/10.1128/microbiolspec.GPP3-0032-2018

Fiedoruk, K., Daniluk, T., Mahillon, J., Leszczynska, K. & Swiecicka, I., 2017. Genetic environment of cry1 genes indicates their common origin. Genome Biol. Evol. 9(9): 2265–2275. https://doi.org/10.1093/gbe/evx165

Francis, K. P., Mayr, R., von Stetten, F., Stewart, G. S. & Scherer, S., 1998. Discrimination of psychrotrophic and mesophilic strains of the *Bacillus cereus* group by pcr targeting of major cold shock protein genes. Appl. Environ. Microbiol. 64(9): 3525–9.

Fu, Y., Wu, Y., Yuan, Y. & Gao, M., 2019. Prevalence and diversity analysis of candidate prophages to provide an understanding on their roles in *Bacillus thuringiensis*. Viruses 11(4): 388. https://doi.org/10.3390/v11040388

Gaidelyte, A., Cvirkaite-Krupovic, V., Daugelavicius, R., Bamford, J. K. & Bamford, D. H., 2006. The entry mechanism of membrane-containing phage Bam35 infecting *Bacillus thuringiensis*. J. Bacteriol. 188(16): 5925–34. https://doi.org/10.1128/JB.00107-06

Garretto, A., Miller-Ensminger, T., Wolfe, A. J. & Putonti, C., 2019. Bacteriophages of the lower urinary tract. Nat. Rev. Urol. 16(7): 422–432. https://doi.org/10.1038/s41585-019-0192-4

Gillis, A., Fayad, N., Makart, L., Bolotin, A., Sorokin, A., et al., 2018. Role of plasmid plasticity and mobile genetic elements in the entomopathogen *Bacillus thuringiensis* serovar *israelensis*. FEMS Microbiol. Rev. 42(6): 829–856. https://doi.org/10.1093/femsre/fuy034

Gillis, A., Guo, S., Bolotin, A., Makart, L., Sorokin, A., et al., 2016. Detection of the cryptic prophage-like molecule pbtic235 in *Bacillus thuringiensis* subsp. *israelensis*. Research in Microbiology 168(4): 319–330. https://doi.org/10.1016/j.resmic.2016.10.004

Gillis, A. & Mahillon, J., 2014a. Phages preying on *Bacillus anthracis, Bacillus cereus*, and *Bacillus thuringiensis*: Past, present and future. Viruses 6(7): 2623–72. https://doi.org/10.3390/v6072623

Gillis, A. & Mahillon, J., 2014b. Prevalence, genetic diversity, and host range of tectiviruses among members of the *Bacillus cereus* group. Appl. Environ. Microbiol. 80(14): 4138–52. https://doi.org/10.1128/AEM.00912-14

Guidi-Rontani, C., Pereira, Y., Ruffie, S., Sirard, J.-C., Weber-Levy, M., et al., 1999. Identification and characterization of a germination operon on the virulence plasmid pxol of *Bacillus anthracis*. Molecular Microbiology 33(2): 407–414. https://doi.org/10.1046/j.1365-2958.1999.01485.x

Gurevich, A., Saveliev, V., Vyahhi, N. & Tesler, G., 2013. Quast: Quality assessment tool for genome assemblies. Bioinformatics, 29(8): 1072–5. https://doi.org/10.1093/bioinformatics/btt086

Houry, A., Briandet, R., Aymerich, S. & Gohar, M., 2010. Involvement of motility and flagella in *Bacillus cereus* biofilm formation. Microbiology 156(4): 1009–1018. https://doi.org/10.1099/mic.0.034827-0

Huerta-Cepas, J., Forslund, K., Coelho, L. P., Szklarczyk, D., Jensen, L. J., et al., 2017. Fast genome-wide functional annotation through orthology assignment by eggnog-mapper. Mol. Biol. Evol. 34(8): 2115–2122. https://doi.org/10.1093/molbev/msx148

Johler, S., Kalbhenn, E. M., Heini, N., Brodmann, P., Gautsch, S., et al., 2018. Enterotoxin production of *Bacillus thuringiensis* isolates from biopesticides, foods, and outbreaks. Front. Microbiol. 9:1915. https://doi.org/10.3389/fmicb.2018.01915

Kalyaanamoorthy, S., Minh, B. Q., Wong, T. K. F., Von Haeseler, A. & Jermiin, L. S., 2017. Modelfinder: Fast model selection for accurate phylogenetic estimates. Nat. Methods 14(6): 587–589. https://doi.org/10.1038/nmeth.4285

Kronstad, J. W., Schnepf, H. E. & Whiteley, H. R., 1983. Diversity of locations for *Bacillus thuringiensis* crystal protein genes. J. Bacteriol. 154(1): 419–428. https://doi.org/10.1128/jb.154.1.419-428.1983

Krupovic, M. & Koonin, E. V., 2015. Polintons: A hotbed of eukaryotic virus, transposon and plasmid evolution. Nat. Rev. Microbiol. 13(2): 105–15. https://doi.org/10.1038/nrmicro3389

Krzywinski, M., Schein, J., Birol, I., Connors, J., Gascoyne, R., et al., 2009. Circos: An information aesthetic for comparative genomics. Genome Res. 19(9): 1639–1645. https://doi.org/10.1101/gr.092759.109

Laurinmaki, P. A., Huiskonen, J. T., Bamford, D. H. & Butcher, S. J., 2005. Membrane proteins modulate the bilayer curvature in the bacterial virus Bam35. Structure, 13(12): 1819–28. https://doi.org/10.1016/j.str.2005.08.020

Letunic, I. & Bork, P., 2016. Interactive tree of life (itol) v3: An online tool for the display and annotation of phylogenetic and other trees. Nucleic Acids Res. 44(W1): W242–5. https://doi.org/10.1093/nar/gkw290

Liu, Y., Du, J., Lai, Q., Zeng, R., Ye, D., et al., 2017. Proposal of nine novel species of the *Bacillus cereus* group. Int. J. Syst. Evol. Microbiol. 67(8): 2499–2508. https://doi.org/10.1099/ijsem.0.001821

Meric, G., Mageiros, L., Pascoe, B., Woodcock, D. J., Mourkas, E., et al., 2018. Lineage-specific plasmid acquisition and the evolution of specialized pathogens in *Bacillus thuringiensis* and the *Bacillus cereus* group. Mol. Ecol. 27(7): 1524–1540. https://doi.org/10.1111/mec.14546

Moumen, B., Nguen-The, C. & Sorokin, A., 2012. Sequence analysis of inducible prophage phis3501 integrated into the haemolysin ii gene of *Bacillus thuringiensis* var *israelensis* atcc35646. Genet. Res. Int. 2012:543286. https://doi.org/10.1155/2012/543286

Nathan, S., Aziz, D. H. & Mahadi, N. M., 2006. Phage displayed *Bacillus thuringiensis* cry1Ba4 toxin is toxic to plutella xylostella. Curr. Microbiol. 53(5): 412–5. https://doi.org/10.1007/s00284-006-0164-9

Nattestad, M., Chin, C.-S. & Schatz, M. C., 2016. Ribbon: Visualizing complex genome alignments and structural variation. bioRxiv. https://doi.org/10.1101/082123

Nguyen, L. T., Schmidt, H. A., von Haeseler, A. & Minh, B. Q., 2015. Iq-tree: A fast and effective stochastic algorithm for estimating maximum-likelihood phylogenies. Mol. Biol. Evol. 32(1): 268–74. https://doi.org/10.1093/molbev/msu300

Page, A. J., Cummins, C. A., Hunt, M., Wong, V. K., Reuter, S., et al., 2015. Roary: Rapid large-scale prokaryote pan genome analysis. Bioinformatics 31(22): 3691–3693. https://doi.org/10.1093/bioinformatics/btv421

Potter, S. C., Luciani, A., Eddy, S. R., Park, Y., Lopez, R., et al., 2018. Hmmer web server: 2018 update. Nucleic Acids Res. 46(W1): W200–W204. https://doi.org/10.1093/nar/gky448

Pruss, B. M., Dietrich, R., Nibler, B., Martlbauer, E. & Scherer, S., 1999a. The hemolytic enterotoxin hbl is broadly distributed among species of the *Bacillus cereus* group. Appl. Environ. Microbiol. 65(12): 5436–42.

Pruss, B. M., Francis, K. P., von Stetten, F. & Scherer, S., 1999b. Correlation of 16s ribosomal DNA signature sequences with temperature-dependent growth rates of mesophilic and psychrotolerant strains of the *Bacillus cereus* group. J Bacteriol. 181(8): 2624–30. https://doi.org/10.1128/JB.181.8.2624-2630.1999

Rambaut, A., 2012. Figtree (version 1.4.0). Available: http://tree.bio.ed.ac.uk/software/figtree/.

Rasko, D. A., Altherr, M. R., Han, C. S. & Ravel, J., 2005. Genomics of the *Bacillus cereus* group of organisms. FEMS Microbiol. Rev. 29(2): 303–329. https://doi.org/10.1016/j.fmrre.2004.12.005

Ravantti, J. J., Gaidelyte, A., Bamford, D. H. & Bamford, J. K., 2003. Comparative analysis of bacterial viruses Bam35, infecting a gram-positive host, and PRD1, infecting gram-negative hosts, demonstrates a viral lineage. Virology 313(2): 401–14. https://doi.org/10.1016/s0042-6822(03)00295-2

Raymond, B. & Bonsall, M. B., 2013. Cooperation and the evolutionary ecology of bacterial virulence: The *Bacillus cereus* group as a novel study system. Bioessays 35(8): 706–16. https://doi.org/10.1002/bies.201300028

Sedlazeck, F. J., Rescheneder, P., Smolka, M., Fang, H., Nattestad, M., et al., 2018. Accurate detection of complex structural variations using single-molecule sequencing. Nat. Methods 15(6): 461–468. https://doi.org/10.1038/s41592-018-0001-7

Siguier, P., 2006. Isfinder: The reference centre for bacterial insertion sequences. Nucleic Acids Res. 34(90001): D32–D36. https://doi.org/10.1093/nar/gkj014

Stajich, J. E., Block, D., Boulez, K., Brenner, S. E., Chervitz, S. A., et al., 2002. The bioperl toolkit: Perl modules for the life sciences. Genome Res. 12(10): 1611–8. https://doi.org/10.1101/gr.361602

Sullivan, M. J., Petty, N. K. & Beatson, S. A., 2011. Easyfig: A genome comparison visualizer. Bioinformatics 27(7): 1009–1010. https://doi.org/10.1093/bioinformatics/btr039

T’Syen, J., Raes, B., Horemans, B., Tassoni, R., Leroy, B., et al., 2018. Catabolism of the groundwater micropollutant 2,6-dichlorobenzamide beyond 2,6-dichlorobenzoate is plasmid encoded in aminobacter sp. Msh1. Appl. Microbiol. Biotechnol. 102(18): 7963–7979. https://doi.org/10.1007/s00253-018-9189-9

Tang, M., Bideshi, D. K., Park, H. W. & Federici, B. A., 2007. Iteron-binding orf157 and ftsz-like orf156 proteins encoded by pbtoxis play a role in its replication in *Bacillus thuringiensis* subsp. israelensis. 189(22): 8053–8058. https://doi.org/10.1128/jb.00908-07

Tatusova, T., DiCuccio, M., Badretdin, A., Chetvernin, V., Nawrocki, E. P., et al., 2016. Ncbi prokaryotic genome annotation pipeline. Nucleic Acids Res. 44(14): 6614–24. https://doi.org/10.1093/nar/gkw569

Verheust, C., Fornelos, N. & Mahillon, J., 2005. GIL16, a new gram-positive tectiviral phage related to the *Bacillus thuringiensis* GIL01 and the *Bacillus cereus* pbclin15 elements. J. Bacteriol. 187(6): 1966–1973. https://doi.org/10.1128/jb.187.6.1966-1973.2005

Wick, R., 2017. Porechop. Available: https://github.com/rrwick/Porechop.

Wick, R. R., Judd, L. M., Gorrie, C. L. & Holt, K. E., 2017. Unicycler: Resolving bacterial genome assemblies from short and long sequencing reads. PLoS Comput. Biol. 13(6): e1005595. https://doi.org/10.1371/journal.pcbi.1005595

Wick, R. R., Schultz, M. B., Zobel, J. & Holt, K. E., 2015. Bandage: Interactive visualization of de novo genome assemblies. Bioinformatics 31(20): 3350–2. https://doi.org/10.1093/bioinformatics/btv383

Zheng, J., Gao, Q., Liu, L., Liu, H., Wang, Y., et al., 2017. Comparative genomics of *Bacillus thuringiensis* reveals a path to specialized exploitation of multiple invertebrate hosts. mBio 8(4). https://doi.org/10.1128/mBio.00822-17

Zheng, J., Guan, Z., Cao, S., Peng, D., Ruan, L., et al., 2015. Plasmids are vectors for redundant chromosomal genes in the *Bacillus cereus* group. BMC Genomics 16(1): 6. https://doi.org/10.1186/s12864-014-1206-5

